# Ecological interactions: Patterns of host utilization by tropical butterflies

**DOI:** 10.1101/2021.12.30.474530

**Authors:** Deepak Naik, Srikrishna Ganaraja Bhat S., Sudeep D. Ghate, M. S. Mustak, R. Shyama Prasad Rao

**Author notes:** Environmental Management and Policy Research Institute, Circle Arch, Vinayaka Nagar, Sahyadri Layout, J.P. Nagar, Bengaluru 560078, India.

## Abstract

Structural complexity of ecological networks facilitate the functional robustness of natural ecosystems. Threatened by the human actions such as habitat destruction and climate change, species may be more or less prone to ecological perturbations depending on the nature of their interactions. We examined the host network of tropical butterflies from the Indian region to see their level of interconnectedness. We manually curated larval host utilization data for 1053 butterflies of India. About 98.8% of species that occur pan-India and 90.6% of species exclusive to the Western Ghats had known hosts whereas it was only 25.9% for species exclusive to north-east India. There were 2589 unique butterfly-host interactions comprising 519 butterfly species and their 1091 known hosts. However, nearly 30% of the species had only single hosts. The Fabaceae and Poaceae were the key host families that accounted for 32.8% of the interactions. There were clear host preferences and monocots hosted disproportionately more butterfly species and interactions. *Vanessa cardui* had at least 39 known hosts while *Ochlandra travancorica* supported 19 butterfly species. There were 2693 species-pairs and 4226 interactions among 469 butterflies due to shared hosts. Many butterfly species that have relatively few/unique hosts might be vulnerable in the context of habitat destruction and climate change. This work has great relevance to the ecology and conservation of butterflies in India.

## Introduction

A very complex web of relationships between species is a key feature which facilitates robustness and resilience of natural ecosystems. Human activities, directly or indirectly, tend to simplify the composition and the structure of ecological networks and removal of a few key components or links can lead to major disruption or irreversible damage to such natural ecosystems (Allesina *et al*., 2009; Landi *et al*., 2018). Currently biodiversity is in the state of decline worldwide due to anthropogenic actions such as agricultural practices, habitat destruction, and climate change (Wepprich *et al*., 2019). Species may be prone to ecological perturbations depending on the nature of their interactions as well as their extent of interconnectedness.

One such important ecological networks is the complex web of interactions between plant feeding (phytophagous) insects such as butterflies and their hosts. Given the presence of myriad plant secondary metabolites, phytophagous insects and their hosts are engaged in a chemical arms race for more than 420 million years (Muto-Fujita*et al*., 2017). As per “oscillation hypothesis”, the availability of a vast diversity of hosts has driven the speciation of phytophagous insects due to divergent selection (Jousselin and Elias, 2019). In fact, interactions between insects and their hosts is central to the diversification of both, and the variability in host use has been shown to drive butterfly diversification (Braga *et al*., 2018). As phytophagous insects make up a large part of the earth’s total biodiversity, exploring their diversification and interactions is central to understanding global biodiversity (Ferrer-Paris *et al*., 2013).

Host utilization by butterflies is a complex process driven by multiple factors such as butterfly choice, and host availability and chemistry (Jones and Agrawal, 2019). Plants serve as nectar hosts for adult butterflies or larval hosts (Curtis et al., 2015; Mukherjee *et al*., 2019; Nitin *et al*., 2018). Distribution and abundance of butterflies is influenced by the availability and abundance of larval host plants and butterflies depend on a narrow set of host plants (Curtis et al., 2015; Nitin *et al*., 2018). For example, Relative abundance of butterflies in island communities is shown to be related to the relative biomasses of their larval host plants (Yamamoto *et al*., 2007). Highly fragmented landscapes due to habitat destruction lead to butterfly meta-populations with complex host-plant dynamics (Hanski and Singer, 2001). Thus, there is an intricate relationship between butterflies and their hosts.

The Western Ghats and North East (Eastern Himalaya and north-eastern India) are key global biodiversity hotspots (Myers *et al*., 2000). While the Western Ghats has over 336 species of butterflies (Nitin *et al*., 2018), the North East has close to 1000 well documented species (Kunte *et al*., 2021). There are numerous endemic, rare, and endangered butterfly species in these regions. While there are many studies to document their diversity, distribution, and abundance (Naik *et al*., 2022; Padhye *et al*., 2012 and numerous references therein), there are only a handful of studies documenting their larval hosts. (Karmakar *et al*., 2018; Kunte *et al*., 2021; Nitin *et al*., 2018). While a few studies have systematically analyzed the larval host utilization patterns for the world’s butterflies (Ferrer-Paris*et al*., 2013; Muto-Fujita *et al*., 2017) such studies are especially scanty for the butterflies of India (Tiple *et al*., 2011).

To fill this gap, we manually curated the host data for the butterflies of India and analysed their interaction network. We discuss the relevance of butterfly-host interaction results in the context of ecology and conservation of butterflies in India.

## Materials and Methods

### Data curation

The larval host data for the butterflies of India were manually curated from the literature (Karmakar *et al*., 2018; Kunte *et al*., 2021; Naik and Mustak, 2020; Naik *et al*., 2022; Nitin *et al*., 2018; Robinson *et al*., 2010; Sengupta *et al*., 2014; Tiple *et al*., 2011; Veenakumari *et al*., 1997, etc.). For example, HOSTS - a database of the world’s lepidopteran host plants (https://www.nhm.ac.uk/our-science/data/hostplants/, last accessed on December 08, 2021) lists host plants for over 380 species of butterflies of India (Robinson *et al*., 2010). Tiple *et al*. (2011) list host plants for 120 species of butterflies of the central India. Nitin *et al*. (2018) list host plants for 320 species of butterflies of the Western Ghats, whereas Karmakar *et al*. (2018) list host plants for 78 species of butterflies of the north-east India. The Butterflies of India website (https://www.ifoundbutterflies.org/, last accessed on December 08, 2021) lists host plants for over 278 butterflies of India (Kunte *et al*., 2021). Altogether, there were over 8,150 entries. However, as there were large overlaps in the host data among different sources, duplicate entries were removed. Further, given the use of different synonyms in different sources, where applicable, latest accepted binomen were used by referring to online literature (http://flora-peninsula-indica.ces.iisc.ac.in/; http://www.worldfloraonline.org/, accessed on December 08, 2021; Borsch *et al*., 2020; Rao *et al*., 2014).

Where available, additional information on the distribution, commonness and abundance, habitat preference, and conservation values of butterflies were collected (Attiwilli et al., 2021; Kunte, 2008; Naik *et al*., 2022; Padhye *et al*., 2012). Similarly, information on the clade and habit of plants were collected from the literature (http://flora-peninsula-indica.ces.iisc.ac.in/; http://www.worldfloraonline.org/, accessed on December 08, 2021; Borsch *et al*., 2020; Rao *et al*., 2014).

### Data/statistical analyses

Overlaps in the number of butterfly species based on the distribution/regions were shown in Venn diagram, and the proportions of species with hosts were computed for each set and shown as pie charts or bar charts. Butterfly-host networks were computed from the unique interactions and shown as incidence matrices at host species, genus, and family-levels. Given that a butterfly species might have multiple hosts and vice versa, we computed the adjacency matrix of first unipartite butterfly-host network (Pavlopoulos *et al*., 2018).

All data handling/analyses were performed in Python; and visualization/graphs were done using R or Microsoft Excel.

Where requisite, a one- or two-proportion Z-test was done to check if the observed proportion was significantly different from the expected. A chi-square test for independence was used to check whether (multiple) sample proportions were significantly different (Agresti, 2007). Statistical tests were done in R.

## Results

### Host utilization by butterflies of India

Of the 1064 listed butterflies of India, 519 species (48.8%) have at least one known host. Individually, 95.5% of the species (n=336) that occur in the Western Ghats (WG), 92.9% (n=367) in Peninsular (PN), 69% (n=150) in Andaman & Nicobar (A&N), and 46.1% (n=934) in the North-East region have known hosts (Fig. 1A, Table S1). The butterflies of the Western Ghats is a subset of peninsular region. Relatively higher proportion of species that occur in two or more regions have known hosts. For example, 98.8% of the species (n=84) that occur pan-India and 96.8% (n=155) that occur in the Western Ghats, Peninsular, and North-East have known hosts. While 90.6% (n=96) of the butterfly species exclusive to peninsular region (which includes the Western Ghats) have known hosts, significantly lower proportion of species (25.9%, n=644, p=1.3E-35, z-test for two proportions) exclusive to North-East have known hosts (Fig. 1B). None of the 32 butterfly species that were exclusive to Andaman & Nicobar have known hosts. A family-wise summary of host plant utilization for different regions is given in Table 1.

**Table 1.** Family-wise and region-wise percentages of butterfly species with known hosts. Numbers in parentheses are sample sizes (n).

| Family | Western Ghats | Peninsular | Andaman & Nicobar | North East | India |
| --- | --- | --- | --- | --- | --- |
| **Hesperiidae** | 97.5 (81) | 94.3 (87) | 91.3 (23) | 42.6 (188) | 48.4 (219) |
| **Lycaenidae** | 89.1 (101) | 87.9 (107) | 81.6 (38) | 45.2 (252) | 46.2 (275) |
| **Nymphalidae** | 99.0 (101) | 94.7 (113) | 48.1 (52) | 42.9 (352) | 45.3 (402) |
| **Papilionidae** | 100.0 (19) | 100.0 (21) | 61.5 (13) | 62.3 (61) | 63.0 (73) |
| **Pieridae** | 96.9 (32) | 94.6 (37) | 77.8 (18) | 61.9 (63) | 63.6 (77) |
| **Riodinidae** | 100.0 (2) | 100.0 (2) | 100.0 (1) | 50.0 (18) | 50.0 (18) |
| **All** | **95.5 (336)** | **92.9 (367)** | **69.0 (145)** | **46.1 (934)** | **48.8 (1064)** |

**Fig. 1.**
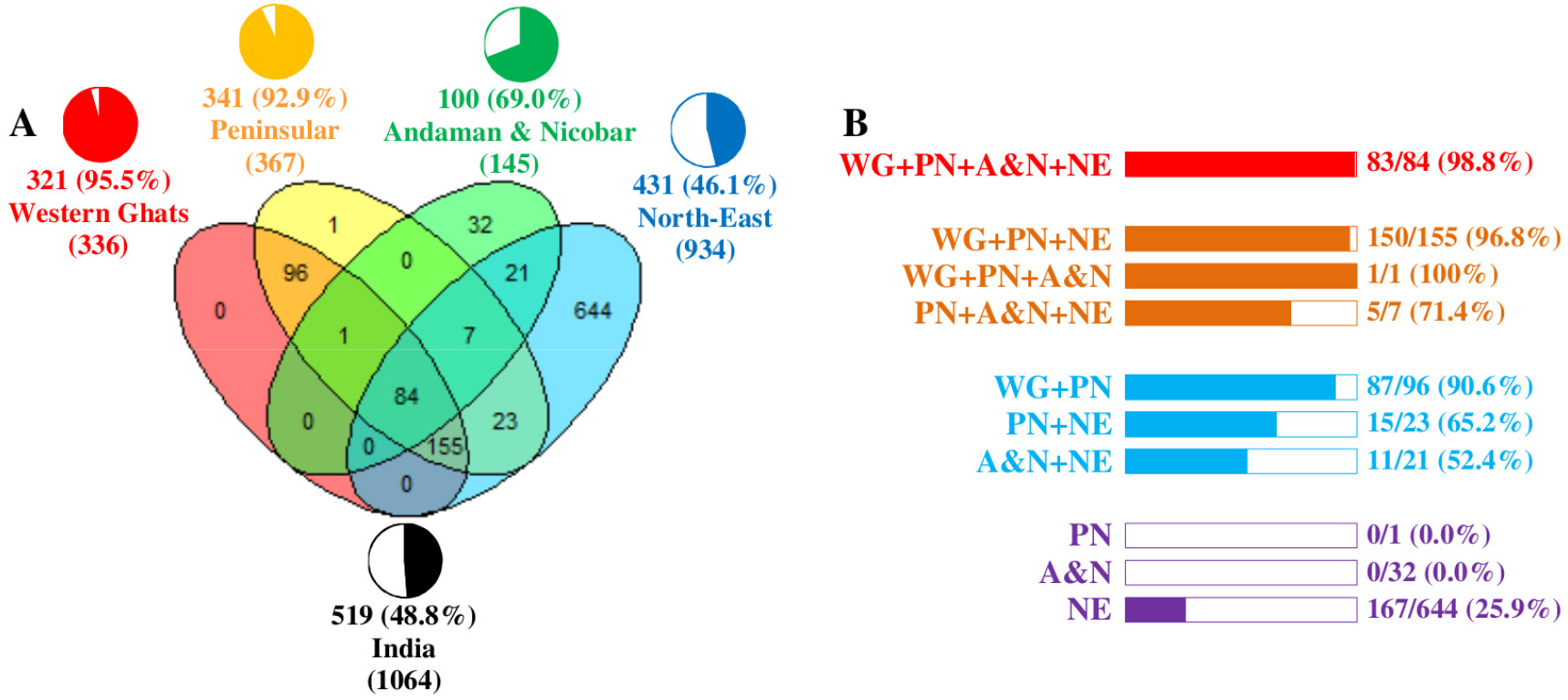
Statistics on host utilization by butterflies of India. (A) Venn diagram shows the overlaps between butterflies of the Western Ghats (WG), Peninsular (PN), Andaman & Nicobar (A&N), and North-East (NE) regions. Of the 1064+ listed butterflies of India, 336 species occur in the Western Ghats whereas 934+ occur in the North-East region. The number and percentage of butterfly species with at least one known host are shown for four regions and for India (pie charts). Nearly 96% of the butterfly species from the Western Ghats have known hosts, whereas it is only 46% for North-East species. (B) About 98.8% of butterfly species that occur pan-India have known hosts. Similarly, relatively higher proportion of species that occur in two or more regions have known hosts. While 90.6% of the butterfly species exclusive to Peninsular region (which includes the Western Ghats) (n=96) have known hosts, just 25.9% of species exclusive to North-East (n=644) have known hosts. Note: The butterflies of the Western Ghats are a subset of the butterflies of the Peninsular region.

### Butterfly-host interaction network

There were 2589 unique butterfly-host interactions comprising 519 butterflies and their 1091 known hosts (Table S1 and S2). While 1024 hosts were known at species-level (from 517 genera), a few others were known only at genus (65) or family (2) levels (Table S2). There were 111 angiosperm host plant families, and a family each from gymnosperm (Cycadaceae) and Insecta (Coccidae). The Fabaceae was the key host family that accounted for 178 (16.3%) hosts and 497 (19.2%) interactions, and supported 101 butterfly species. The next key family Poaceae accounted for 77 (7.1%) hosts and 353 (13.6%) interactions, and supported 116 butterfly species. The monocot clade which contained 172 hosts in 14 families supported 157 butterfly species, while the non-monocot clade (Eudicots and Magnoliids) with 914 hosts supported 381 butterfly species. For the given number of hosts, the monocot clade had significantly more interactions with butterfly species compared to non-monocots (561 interactions for 172 hosts versus 2022 for 914 respectively, Table S3, p=7.6E-5, chi-square test).

Numerous butterfly species had multiple interactions with the hosts (at family-, genus-, and species-levels) and vice versa (Fig. 2). For example, *Vanessa cardui* had at least 39 known hosts each of which in turn supported many other butterfly species. Similarly, *Ochlandra travancorica* supported 19 butterfly species which in turn had many other hosts. However, there was no interaction between *Vanessa cardui* and *Ochlandra travancorica*, and none of hosts of *Vanessa cardui* and the butterfly species supported by *Ochlandra travancorica* had any interactions with each other thus making it an empty bipartite graph (Fig. 3). There was a divide between butterfly species in using either monocot or non-monocot species. For example, Fabaceae supported 101 species of butterflies while Poaceae supported 116. Just two species - *Zizula hylax* and *Pelopidas mathias* fed on hosts belonging to both the families (Fig. S1A). While monocots supported 157 species of butterflies, non-monocots supported 382. Twenty species fed on hosts from both the clades. However, majority of them had significantly more hosts (p<0.05, z-test for one proportion) from one of the clades (Fig. S1B). About 47.7% of butterfly species with at least two hosts (n=365) interacted with diverse hosts from two or more families. For example, *Rapala manea* had 22 hosts from as many as 14 families including Fabaceae and Phyllanthaceae. *Vanessa cardui* had 39 hosts from six families including Asteraceae and Fabaceae. On the other hand, some species, for example, *Eurema hecabe* and *Lampides boeticus* while fed on 28 and 26 hosts respectively, were restricted to just one family Fabaceae.

**Fig. 2.**
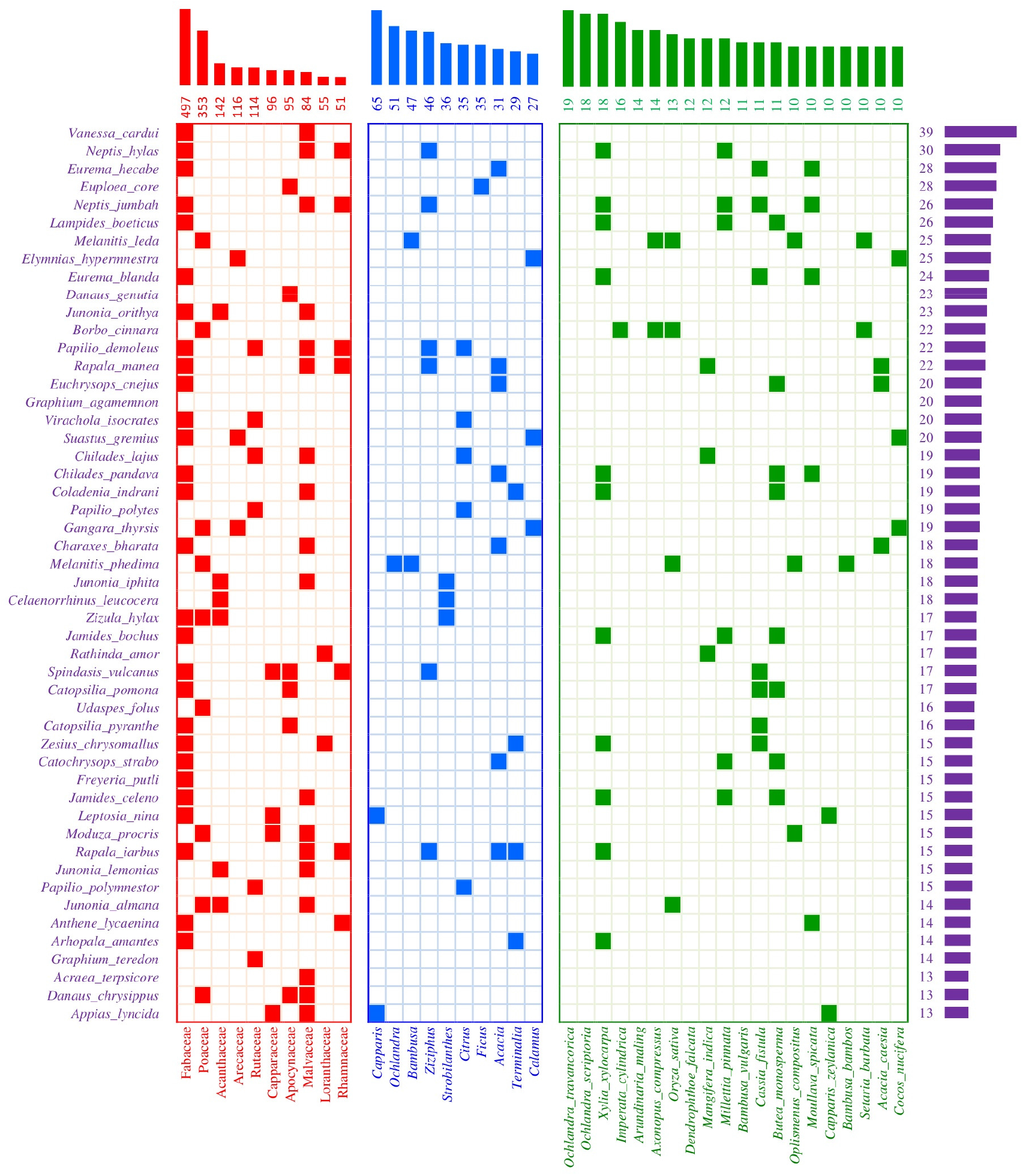
Butterfly-host network. Incidence matrices show the interactions (filled cells) of the top 50 butterfly species (which represent 37% of total interactions) with (A) top 10 host families (which represent 61.9% of total interactions), (B) top 10 host genera (which represent 15.7% of total interactions), and (C) top 20 host species (which represent 10.8% of total interactions). The bar graphs show the total number of interactions. Fabaceae is the top host family with 497 interactions (from 101 butterfly species and 178 hosts). While *Vanessa cardui* is the top butterfly species, none of the top host genera and species support it. Similarly, *Ochlandra travancorica* and *Ochlandra scriptoria* are the top hosts, but none of the top butterfly species feed on them. Lists are sorted in descending order of total number of interactions.

**Fig. 3.**
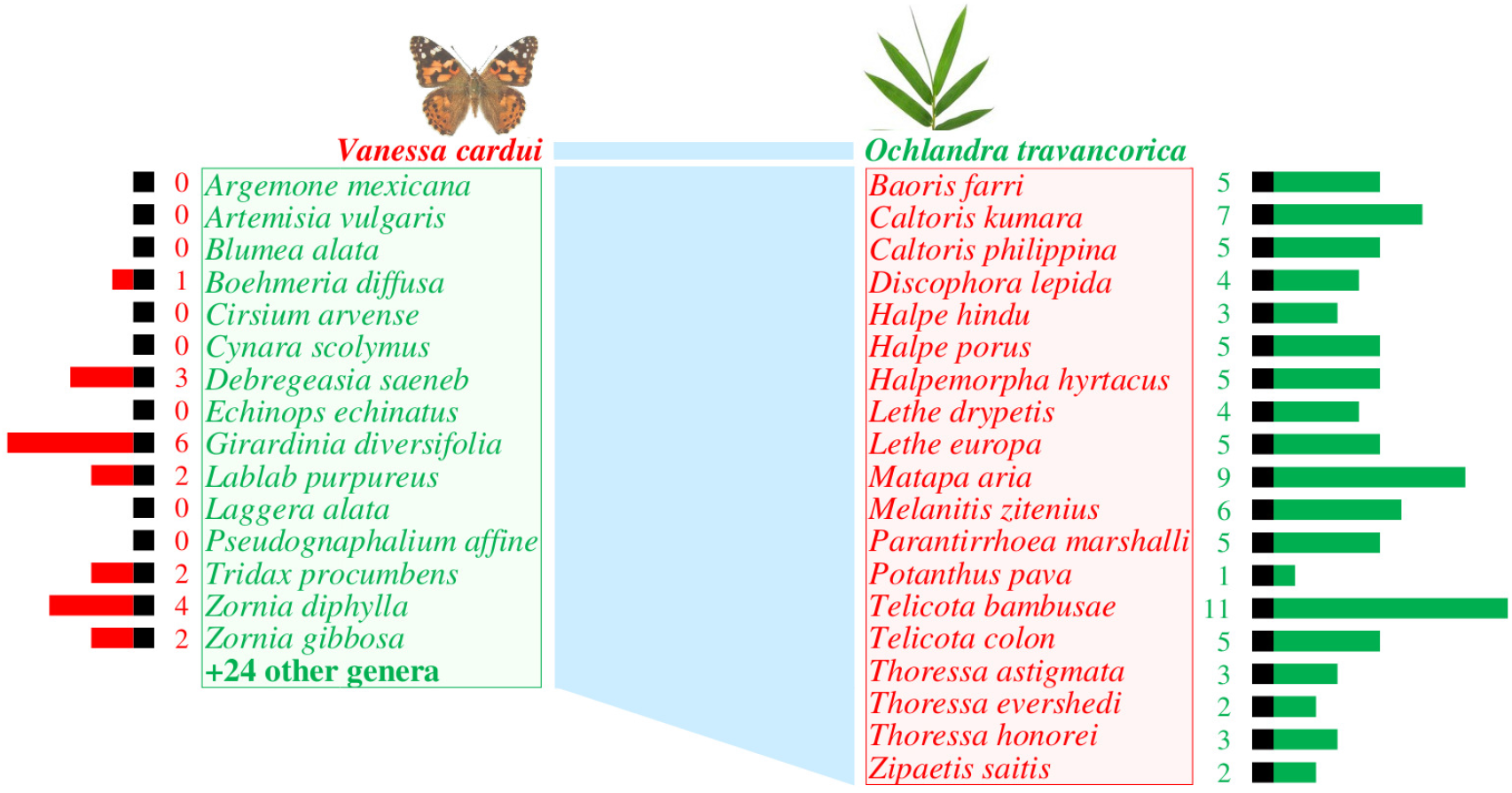
Interactions of the top species. *Vanessa cardui* has at least 39 known hosts (green box, hosts identified only to the genus level are not listed) whereas *Ochlandra travancorica* is a host to 19 butterfly species (red box). Each of the 39 hosts in turn support numerous other butterfly species (red bars and numbers). Similarly, each of the 19 butterfly species also have numerous other hosts (green bars and numbers). An empty bipartite graph of these two sets of species is shown at the center as light blue shade - *Ochlandra travancorica* is not a host for *Vanessa cardui* and none of the species listed interact with each other. Species are sorted in alphabetical order.

### Butterfly-butterfly interactions

The adjacency matrix of first unipartite butterfly-host network is given in Table S4. There were as many as 2693 species-pairs (4226 interactions) among 469 butterflies due to shared hosts. About 30.2% of those species-pairs had two or more shared hosts (2346 interactions). There were 212 species-pairs (550 interactions) among top 50 butterfly species based on the number of hosts (Fig. 4A). About 54.2% of those pairs had two or more shared hosts. For example, *Jamides bochus* has 17 hosts (all from Fabaceae) and *Lampides boeticus* has 26 hosts (all from Fabaceae) with 12 hosts in common (Fig. 4B). Overall, *Neptis jumbah* (26 hosts) had 102 interactions with 62 species. A similar looking *Athyma perius* (13 hosts) had just seven interactions with three species. *Rapala manea* (22 hosts from 14 families) had 78 interactions with 52 species. Although *Vanessa cardui* has 39 hosts, it had modest 25 interactions with 17 species. In comparison, *Caltoris kumara* (8 hosts) had 86 interactions with 44 species due to general hosts such as *Ochlandra travancorica, Bambusa vulgaris*, and *Oryza sativa*. On the other hand, some species such as *Burara gomata* (8 hosts) and *Papilio machaon* (7 hosts) had no interactions with others (Table S4).

**Fig. 4.**
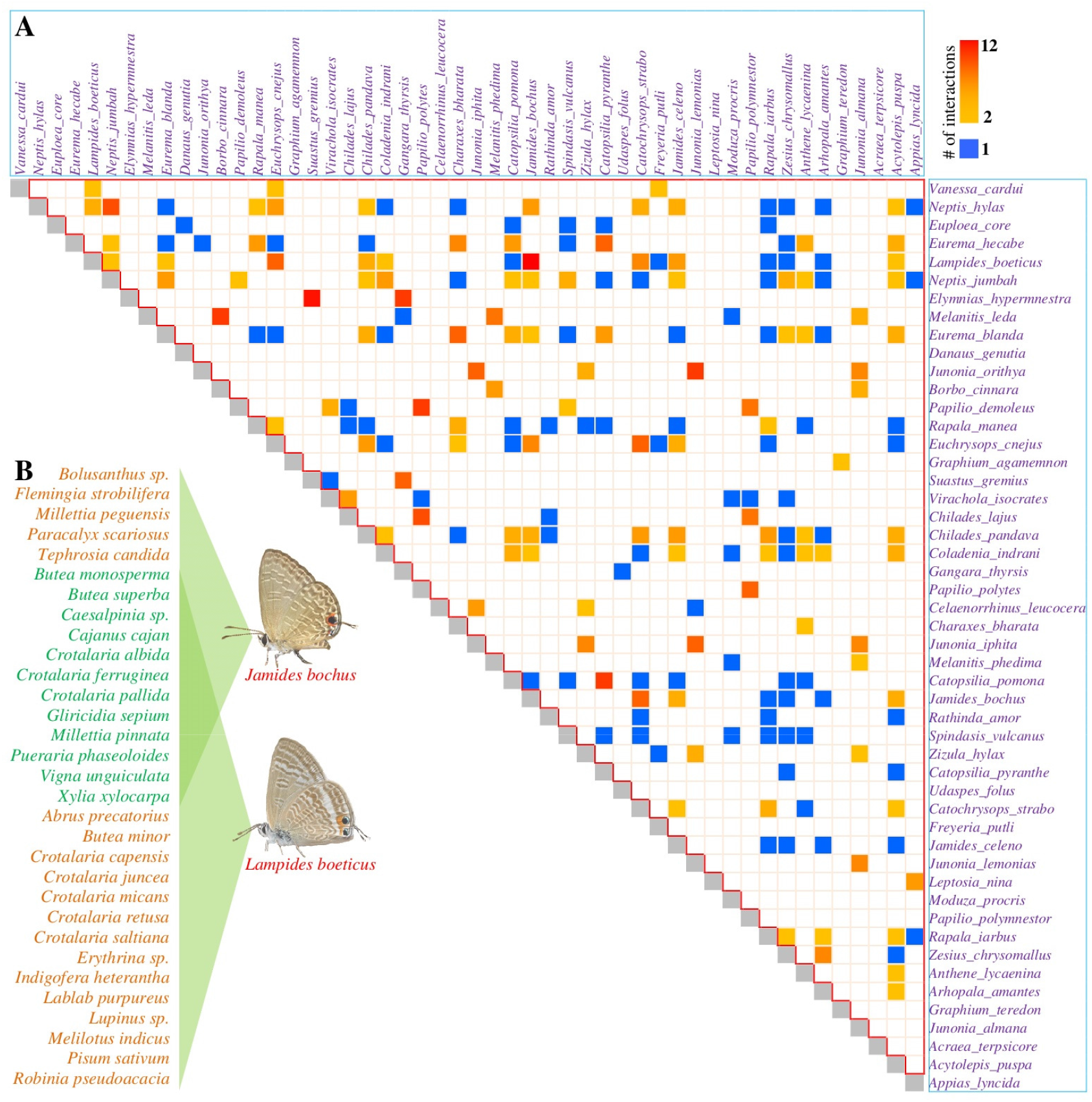
Adjacency matrix of first unipartite butterfly-host network. Overall, there were 2693 interacting species-pairs (4226 interactions among 469 butterflies) with 30.2% of species-pairs with two or more shared hosts and 2346 interactions (see Table S4). (A) The matrix shows interactions among the top 50 butterfly species based on the number of hosts. Here, 54.2% of the interacting species-pairs (n=212) had two or more shared hosts. For example, (B) *Jamides bochus* has 17 hosts (all from Fabaceae) and *Lampides boeticus* has 26 hosts (all from Fabaceae) with 12 hosts in common.

## Discussion

The Western Ghats and North East regions of India have a very rich diversity of butterflies. However, their ecology and host plants in particular are poorly explored, and only patchy information is available on their hosts (Karmakar *et al*., 2018; Kunte *et al*., 2021; Naik and Mustak, 2020; Naik *et al*., 2022; Nitin *et al*., 2018; Robinson *et al*., 2010; Sengupta *et al*., 2014; Tiple *et al*., 2011; Veenakumari *et al*., 1997; etc.). In this work, we provided 2589 unique butterfly-host interactions comprising 519 butterflies and their 1091 known hosts. Yet, this is tiny compared to 3435 interactions known for Japanese butterflies (Muto-Fujita*et al*., 2017) and 44118 records for 5152 species of butterflies worldwide (Ferrer-Paris *et al*., 2013). The HOSTS database has over 102981 records for 20457 lepidopteran species (butterflies and moths), but contains only 1185 records for 380 species of butterflies of India (Robinson *et al*., 2010). The Butterflies of India website has information on larval hosts for over 278 species (Kunte *et al*., 2021).

Hosts for nearly all pan-Indian butterfly species are known as there is a better chance of encountering them in the field. In contrast, hosts of butterflies exclusive to only one region are poorly known. This is particularly stark for butterflies of North East and Andaman & Nicobar (Veenakumari *et al*., 1997). Thus, there is a need to explore the hosts of butterflies of these regions in the context of ecology and conservation (Karmakar *et al*., 2018).

There were clear host preferences and monocots hosted disproportionately more butterfly species and interactions. The Fabaceae and Poaceae are known to be preferred host families (Ferrer-Paris *et al*., 2013; Muto-Fujita *et al*., 2017; Tiple *et al*., 2011), and associations between certain groups of butterflies and hosts such as between Papilionidae and magnoliids, and Hesperiidae and monocots are recognized (Ferrer-Paris *et al*., 2013). Numerous factors drive the host utilization by butterflies, including distribution and availability of hosts, female butterfly’s oviposition preference, larval performance, and the presence of competition (Jones and Agrawal, 2019). In addition, the presence of photochemicals and metabolic processes play a key role in host selection (Muto-Fujita *et al*., 2017). One key family of plants that was hardly used as host is Liliaceae and this is possibly due to its unique phytochemistry (Li *et al*., 2006).

Many butterfly species have multiple hosts and vice versa. For example, *Vanessa cardui* has at least 39 hosts in India, and as the most widespread species it is known to have over 300 hosts worldwide (Robinson *et al*., 2010). Host usage might be fluid as ongoing climate-driven alteration in hosts is known in European butterflies (Braschler and Hill, 2007; Pateman *et al*., 2012). Conversely, Muto-Fujita *et al*. (2017) showed the multiple utilization of same hosts by butterflies of different families and demonstrate the independent acquisition of adaptive phenotypes to the same hosts, which could partly due to shared chemistry. Butterfly species that use numerous hosts might still be vulnerable to ecological perturbations depending on preference or functional interactions with their hosts (Allesina *et al*., 2009).

It is known that adaptation to feeding on different hosts can lead to ecological specialization of populations and subsequent speciation (Braga *et al*., 2018; Jousselin and Elias, 2019). While habitat fragmentations might hasten this process, they are also likely to hasten species decline and loss due to drift and ecological perturbations. This effect might be even more drastic in butterfly species that depend on a few/unique hosts. As butterflies are declining due to increasing anthropogenic challenges (Rao and Girish, 2007; Wepprich *et al*., 2019), there is a need to explore and conserve butterfly hosts and their habitats. The quality of habitat patches for specialist species is shown to depend on the host resource (Curtis et al., 2015). Fortunately, a large number of cultivated plant species in the urban areas are known to serve as butterfly hosts (Tiple *et al*., 2011).

To comment on a key limitation of this study, as there were no substantial region-specific data, differences, if any, in the host-plant usage pattern among common butterflies in different geographic regions were not explored. Although data were extensively cleaned, due to taxonomic updates and numerous synonyms, there might still be some inadvertent inaccuracies or overlaps among plant species.

In conclusion, we showed that only 25.9% of the butterfly species exclusive to north-east India have known hosts. We provided curated information of over 2589 unique host interactions for 519 species of butterflies of India. Nearly a third of them had only single known hosts. There were clear host preferences and monocots hosted disproportionately more butterfly species and interactions. There were 2693 species-pairs and 4226 interactions among 469 butterflies due to shared hosts. Given the pace of habitat destruction and climate change, there is a great need to explore the larval hosts in the context of ecology and conservation of butterflies in India.

## Funding and Acknowledgments

This work did not receive any specific funding.

## Statement of Ethics

The work is in compliance with ethical standards. No ethical clearance was necessary.

## Conflict of Interest

The authors declare that there is no conflict of interest.

## Data Availability

The data used in this work were collected/curated from the literature. Relevant data are given in supplemental Tables S1, S2, and S4. Further data may be obtained from the authors for collaborative work upon reasonable request.

## Author Contributions

DN and RSPR initiated the work. DN curated the data. DN and RSPR analyzed the data and wrote the paper. All authors contributed and were involved in the revision.

## Supplemental Information

Supplemental information for this article is available online.

## Notes

### Competing Interest Statement

The authors have declared no competing interest.

