## Supplemental for "Ecological interactions: Patterns of host utilization by tropical butterflies"

[1]#2-2-25-44D, Laxmishreenidhi, Pambattamajal, Kabaka Village, Puttur 574203, India
[2]Vivekananda College, Nehru Nagar, Puttur 574203, India
[3]Nitte University Centre for Science Education and Research (NUCSER), NITTE University, Mangaluru 575018, India
[4]Department of Applied Zoology, Mangalore University, Konaje 574199, India
[5]Biostatistics and Bioinformatics Division, Yenepoya Research Center, Yenepoya University, Mangaluru 575018, India

[+]Current address
Environmental Management and Policy Research Institute, Circle Arch, Vinayaka Nagar, Sahyadri Layout, J.P. Nagar, Bengaluru 560078, India

*Corresponding authors



Running title: Larval hosts of Indian butterflies

*Keywords: Butterfly conservation, Eco-informatics, Ecological interaction network, Public awareness and outreach, Species loss, Systems ecology, Urban biodiversity*

**Table S1.** List of butterfly species, their distribution within India, and the number of known host interactions (Table_S1.xlsx).

**Table S2.** List of known hosts for the butterflies of India (Table_S2.xlsx).

**Table S3.** Clade-wise number of hosts and interactions.

| | # of hosts | # of interactions |
|---|---|---|
| **Eudicots** | 872 | 1938 |
| **Magnoliids** | 42 | 84 |
| **Monocots** | 172 | 561 |
| **Gymnosperms** | 4 | 4 |
| **Insecta** | 1 | 2 |
| **All** | **1091** | **2589** |

**Table S4.** Adjacency matrix of first unipartite butterfly-host network (Table_S4.xlsx).

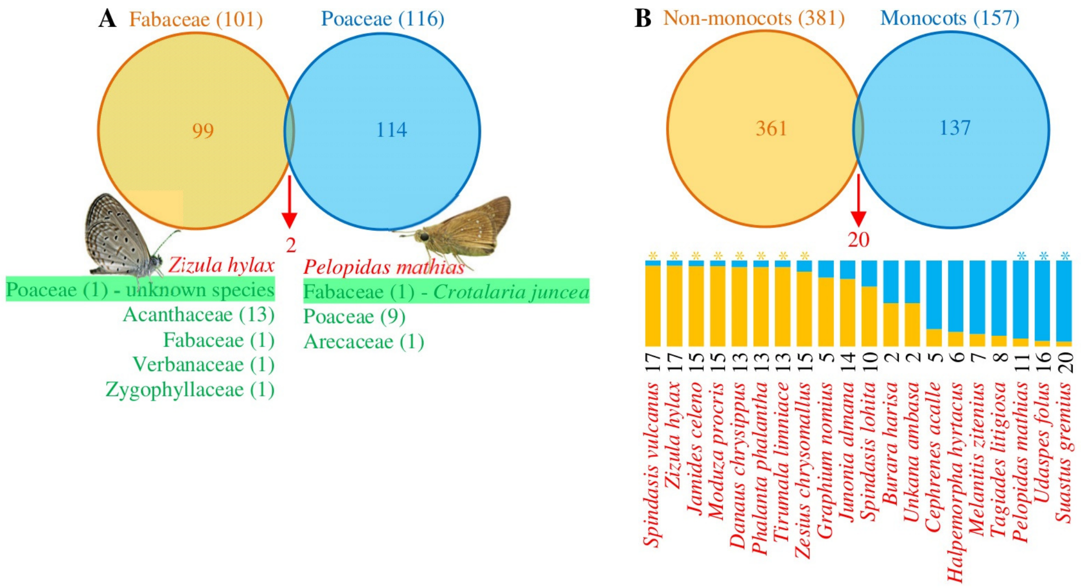


**Fig. S1.** Monocot divide. (A) Fabaceae supports 101 species of butterflies while Poaceae supports 116. Just two species - *Zizula hylax* and *Pelopidas mathias* feed on hosts belonging to both the families. However, those overlaps might be doubtful as *Zizula hylax* mainly feeds on multiple hosts in Acanthaceae and other Eudicots, while its single host in Poaceae was not identified (Robinson *et al*., 2010). Similarly, *Pelopidas mathias* mainly feeds on multiple hosts in Poaceae. (B) While monocots support 157 species of butterflies, non-monocots support 381. Twenty species feed on hosts from both the clades. However, majority of them have significantly more hosts (* indicates $p<0.05$, z-test for one proportion) from one of the clades (stacked bar graph and number of hosts).
